# CellReasoner: A reasoning-enhanced large language model for cell type annotation

**DOI:** 10.1101/2025.05.20.655112

**Authors:** Guangshuo Cao, Yi Shen, Jianghong Wu, Haoyu Chao, Ming Chen, Dijun Chen

## Abstract

We present CellReasoner, a lightweight, open-source large language model (LLM) tailored for single-cell type annotation. We introduced a compact training strategy that activates the reasoning capabilities of 7B-parameter LLMs using only 380 high-quality chain-of-thought exemplars. CellReasoner directly maps cell-level gene expression profiles to cell type labels, exhibiting robust zero- and few-shot generalization. The model further demonstrates expert-level, marker-by-marker reasoning, enabling structured, interpretable annotations and offering a practical solution for intelligent single-cell analysis.

## Main

Cell type annotation is a fundamental step in single-cell data analysis, and it also represents a reasoning process that integrates diverse sources of evidence — such as gene expression profiles, canonical marker genes, and reference datasets — to accurately infer cellular identities. Similar to step-wise inference in artificial intelligence, this process relies on integrating prior knowledge with context-specific features to support high-confidence classification. Recent advances in large language models (LLMs) have demonstrated that, once scaled to sufficient parameter sizes, these models exhibit sophisticated reasoning capabilities across mathematical, logical, and programming tasks ^1,2,3,4^ . This progress highlights the potential for leveraging LLM-based reasoning paradigms to enhance complex biological inference tasks such as automated cell type annotation.

Although several single-cell–specific models ^5,6,7,8^ trained on large datasets have recently emerged, they largely depend on data-intensive paradigms that incur substantial computational costs. Existing annotation workflows often rely on manual marker curation or reference-based mapping, although recent approaches have begun exploring general purpose LLMs (e.g., GPT-4^9^) as automated alternatives^10^. However, these models suffer from limited interpretability and are challenging to deploy or reproduce due to their massive scale, dependency on proprietary APIs, and associated cost and privacy concerns. To overcome these limitations, we developed CellReasoner, a lightweight LLM tailored for cell type annotation based on single-cell transcriptomic data and efficiently deployable on consumer-grade GPUs. Inspired by recent findings that LLMs encode latent biological knowledge^11^, we evaluated open-source LLMs such as Qwen2.5-7B-Instruct^12^ and found that even at a modest scale, they capture rich marker-cell type associations (**Supplementary Fig. 1a,b**). These observations shift the core challenge from knowledge acquisition to knowledge activation — i.e., devising effective strategies to elicit, contextualize, and apply the embedded biological knowledge to real-world annotation tasks.

As a proof of concept, we compiled a benchmark dataset comprising 37,187 cells — referred to as Pancancer38k — aggregated from multiple published pan-cancer single-cell studies (**Supplementary Table 1**). This dataset encompasses diverse tumor types and tissue origins, providing a challenging yet representative testbed for evaluating the annotation accuracy, generalizability, and interpretability of CellReasoner across heterogeneous biological contexts. To serve as model input, we generated natural language-like cell representations (“cell sentences”) based on ranked highly variable genes (HVGs), selected from millions of single-cell profiles in Pancancer38k (**Fig. 1a,b** and see **Methods**). To activate the reasoning capabilities of the model, we developed a novel three-stage training strategy, termed *Cell Reasoning and Annotation Fusion Training* (CRAFT). In the first stage (reasoning scaffold), a small number of chain-of-thought (CoT) exemplars are used to effectively elicit step-wise reasoning in LLMs. The second stage (knowledge infusion) aims to integrate both accurate reasoning and direct annotation, enabling the model to consolidate biological knowledge with practical annotation skills. Finally, the third stage (reasoning mode fusion) addresses the decline in reasoning ability observed during the knowledge infusion stage (**Fig. 1c**).

**Figure 1.**
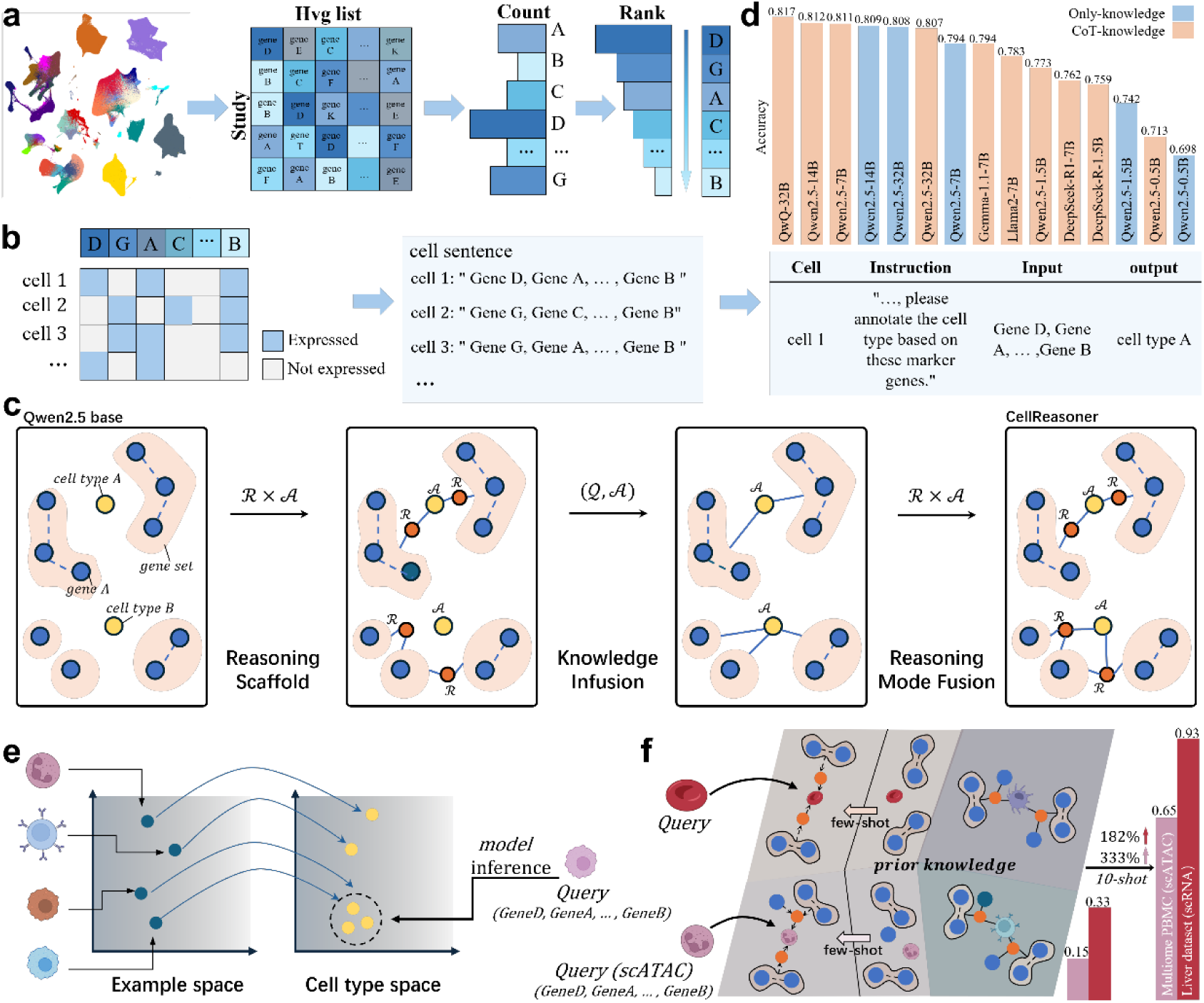
Overview of CellReasoner and the CRAFT training strategy. **a**, Global HVG list construction from 570 studies based on frequency ranking. **b**, Cell sentence generation by encoding expressed genes in a fixed HVG order. **c**, The CRAFT framework includes three stages: reasoning scaffold, knowledge infusion, and reasoning mode fusion. **d**, Accuracy comparison across LLMs of varying parameter sizes. Models trained with both reasoning scaffold and knowledge infusion (CoT-knowledge, orange) outperform those trained with knowledge infusion only (only-knowledge, blue), highlighting the benefit of CoT-based reasoning activation. **e**, Zero-shot application using direct cell-level inputs without clustering. **f**, Few-shot generalization across modalities and tissues. In 10-shot settings, CellReasoner improves accuracy by 182% on scRNA-seq and 333% on scATAC-seq.

CRAFT leverages only 380 high-quality CoT exemplars to elicit step-wise reasoning in open-source LLMs such as Qwen and LLaMA^13^ across varying parameter scales. Model performance was evaluated on an independent test set comprising 3,800 previously unseen cells. Notably, introducing a small number of CoT exemplars during the reasoning scaffold stage markedly improved annotation accuracy compared to the knowledge infusion stage alone (**Fig. 1d**), highlighting the critical role of explicit reasoning guidance in training. Among the models tested, Qwen2.5-7B-Instruct outperformed larger variants (14B and 32B) while incurring lower computational cost, highlighting superior parameter efficiency. Only Qwen-family models surpassed 0.8 accuracy; accordingly, we used Qwen2.5-7B-Instruct and QwQ-32B-Instruct as the foundation models for CellReasoner in the subsequent development and evaluation experiments.

Through limited exemplar learning, CellReasoner acquired the ability to map cell sentences directly to cell types without manual labels or reference databases, enabling robust zero-shot inference (**Fig. 1e**). The model outputs cell type predictions directly, requiring no additional filtering or post-processing, thereby offering a high degree of automation and practical utility (**Supplementary Fig. 1c**). Building on prior biological knowledge, CellReasoner is compatible not only with cell sentences derived from scRNA-seq expression matrices but also with those generated from scATAC-seq gene activity matrices. Under few-shot settings, the model rapidly adapts to new data distributions, enabling cross-modality transfer (**Fig. 1f**).

We evaluated CellReasoner in a zero-shot setting on a pancreatic ductal adenocarcinomas (PDAC) dataset^15^ using cell sentences constructed from top-k ranked genes. Annotation performance improved with increasing sentence length, reaching a plateau at k = 1,000-1,500 for CellReasoner-7B. Similarly, CellReasoner-32B achieved its peak accuracy of 0.73 at k = 1,000 (**Fig. 2a**). A recent study has shown that ChatGPT exhibits some capability in cell type annotation; however, due to its high cost and limited interpretability, it has been more commonly applied to marker gene annotation at the cluster level, following prior clustering. For comparisons, we benchmarked models by providing each cell cluster’s top marker genes as input. Even in this simplified setting, CellReasoner outperformed general-purpose models such as ChatGPT and DeepSeek in both accuracy and robustness (**Fig. 2b**). Further analysis revealed CellReasoner provided finer-grained subtype resolution in the PDAC dataset. For instance, ChatGPT-4o misclassified all CD4⁺ T cell subtypes as CD8⁺ T cells, while DeepSeek-R1 broadly labeled both as “cytotoxic T cells” (**Supplementary Fig. S2a**). To mitigate label imbalance, we constructed a class-balanced test set by sampling 20 cells per type. In this setting, CellReasoner-7B and 32B achieved average scores of 0.72 and 0.75, respectively — substantially outperforming DeepSeek-V3-671B^16^ (0.46) and DeepSeek-R1-671B (0.52) — further confirming CellReasoner’s stability and generalizability (**Fig. 2c**).

**Figure 2.**
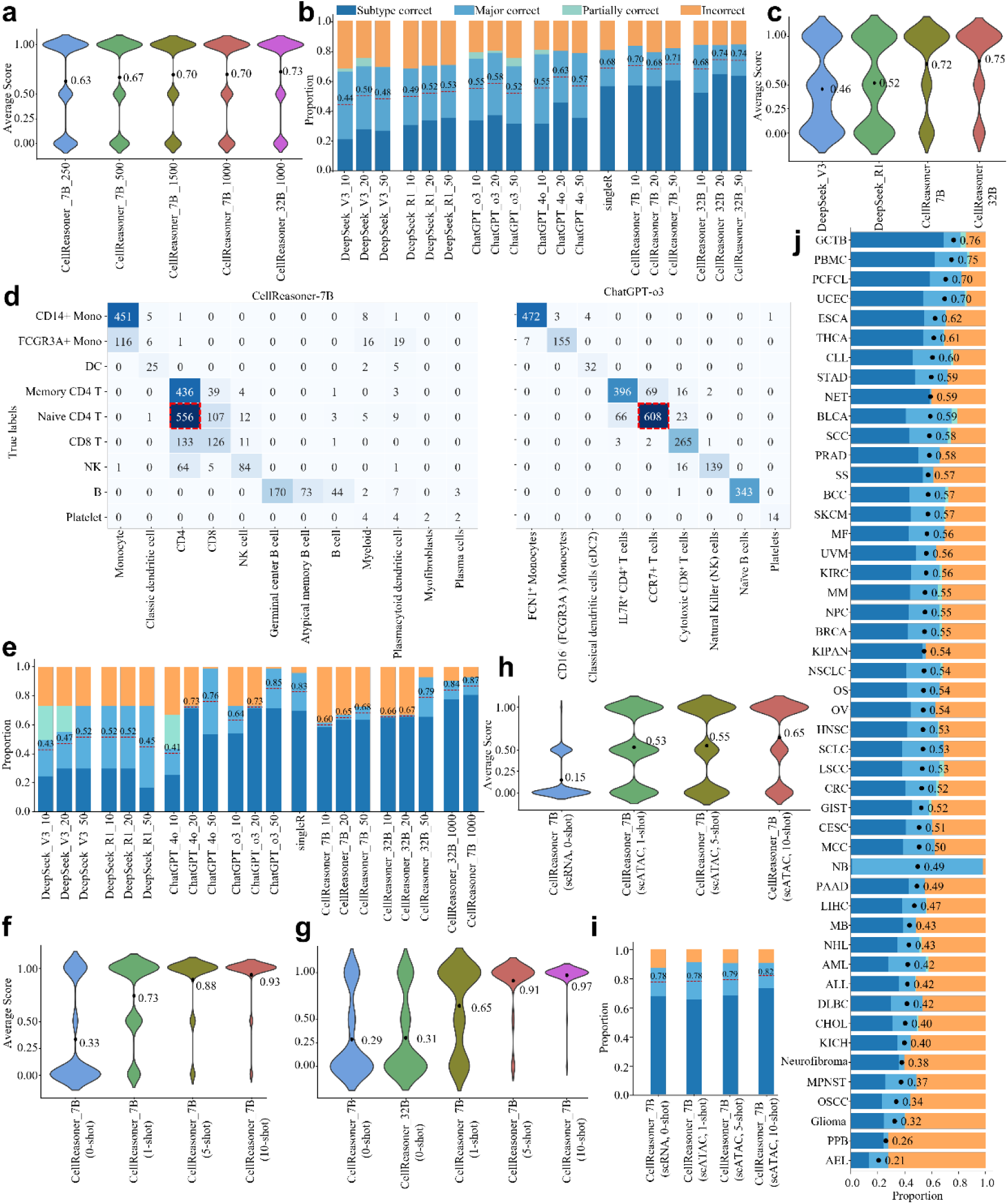
Benchmarking CellReasoner on diverse annotation scenarios. **a**, Zero-shot performance on the PDAC dataset using cell sentences constructed from top-k globally ranked HVGs. **b**, Cluster-level annotation performance on the PDAC dataset using top-k marker genes. Bars show proportions of Subtype correct, Major correct, Partially correct, and Incorrect predictions (top 10/20/50 markers). **c**, Average annotation score on a balanced PDAC subset with 20 cells per cell type. **d**, Confusion matrix comparison on the PBMC3K dataset. Left: CellReasoner-7B with top-1000 HVGs; Right: ChatGPT-o3 with top-50 markers. **e**, Annotation performance on PBMC3K using markers (10/20/50) and HVG-based cell sentences (1000). CellReasoner-7B achieves the highest performance. **f**, Few-shot performance on the liver dataset. **g**, Few-shot performance on a class-balanced subset of the liver dataset. **h**, Few-shot generalization from scRNA-seq to scATAC-seq on the multiome PBMC dataset. **i**, Few-shot generalization to the paired scRNA-seq modality from the multiome PBMC dataset. **j**, Large-scale evaluation across 181 datasets spanning 48 cancer types

We next evaluated CellReasoner on cross-tissue generalization using the PBMC3K benchmark dataset, which is derived from non-cancer peripheral blood samples. CellReasoner-7B correctly annotated both naive CD4+T cells and memory CD4+T cells as CD4+T cells (**Fig. 2d**). In contrast, ChatGPT-o3 misclassified most naive CD4⁺ subsets as generic “T cells,” likely due to its reliance on cluster-level marker gene input. While this approach is simpler and cost-effective, it is vulnerable to error propagation: when clustering is imperfect, a single mislabel can affect all cells within the specific cluster. By comparison, CellReasoner performs cell-type inference at the single-cell level, enabling fine-grained predictions and local error correction. Interestingly, ChatGPT-o3 predicted FCGR3A⁺ monocytes with the exact marker name recall, suggesting that PBMC3K data may have been present in its pretraining corpus. Further comparisons showed CellReasoner consistently outperformed general models like ChatGPT and DeepSeek in subtype-level annotation. Notably, CellReasoner-7B achieved the highest score using the top 1,000 ranked genes (**Fig. 2e**), while CellReasoner-7B exceeded its larger counterpart under class-balanced evaluation (**Supplementary Fig. 2b**).

To evaluate out-of-distribution generalization, we used a liver dataset^14^, which includes cell types absent from training — such as hepatic stellate cells and γδT cells. In the zero-shot setting, CellReasoner-7B was only able to correctly identify previously seen types such as NK and plasma cells (**Supplementary Fig. 2c**), yielding an average prediction score of 0.33. Remarkably, 1-shot fine-tuning boosted the average score to 0.73 (**Fig. 2f**), demonstrating the capacity of the CellReasoner model to rapidly adapt and activate reasoning mechanisms for previously unseen cell types. Accuracy further improved with 5-shot and 10-shot inputs, reaching 0.93 on the full test set (**Supplementary Fig. 2d**) and 0.97 on a class-balanced subset (**Fig. 2g**). In this regard, 10-shot fine-tuning resulted in a 333% increase in accuracy compared to the zero-shot setting. The above results highlight CellReasoner’s robust generalization to novel cell types, efficient knowledge integration, and rapid adaptability, which is crucial for real-world scenarios involving tissue-specific or incompletely annotated data.

To assess CellReasoner’s cross-modality transferability, we conducted few-shot evaluations using multiome peripheral blood mononuclear cell (PBMC) datasets containing both scRNA-seq and scATAC-seq profiles. Although the model was trained solely on scRNA-seq data, we tested its ability to generalize to scATAC-seq inputs by constructing cell sentences from gene activity matrices. In the zero-shot setting, performance was limited (**Fig. 2h**). Remarkably, without any architectural modifications, 1-shot fine-tuning raised the average score to 0.53, and 10-shot input further improved it to 0.65 (182% improvement over zero-shot; **Fig. 1f**) — demonstrating robust adaptability across modalities (**Supplementary Fig. 3a**). Importantly, fine-tuning on scATAC-seq data did not compromise the model’s performance on the original scRNA modality. On the contrary, evaluation on the matched scRNA-seq test set showed consistent or slightly enhanced accuracy as support examples increased (**Fig. 2i** and **Supplementary Fig. 3b**), underscoring the model’s robustness and retention of previously learned knowledge. These results indicate that CellReasoner can acquire modality-specific knowledge while preserving prior understanding, supporting both effective transfer and conservative generalization.

To evaluate CellReasoner’s scalability and robustness across diverse cancer contexts, we tested it on 181 public single-cell datasets spanning 48 cancer types, curated from the TISCH2 database^17^ (**Fig. 2j** and **Supplementary Fig. 3c**). For each dataset, 100 cells per type were randomly sampled to create class-balanced test sets and reduce bias. The model achieved consistently high scores across most datasets, indicating strong cross-tissue adaptability. However, this overall performance does not imply uniform success across all datasets. For instance, in multiple myeloma (MM), CellReasoner achieved a high accuracy of 0.95 on the GSE154763 dataset but only 0.48 on GSE161801 (**Supplementary Fig. 3c**). Such variability likely reflects differences in subtype annotation granularity or the presence of biologically complex or highly specific populations. These results suggest that generalization depends not only on tissue origin but also on the consistency and standardization of cell type definitions across datasets.

To assess whether CellReasoner exhibits expert-level reasoning, we provided a cell sentence derived from a CD4⁺ T cell and asked the model to infer the label based solely on marker genes (**Supplementary Fig. 1d**). CellReasoner-7B correctly identified CD3 and CD4/CD8 as components of the TCR complex, linked IL7R to memory T cell activation, and applied negative exclusion and context-aware inference to predict the correct CD4⁺ T cell identity. In contrast, Qwen2.5-7B produced repetitive CD markers without reaching a conclusion, while DeepSeek-R1-671B, though partially logical, yielded vague outputs like “tumor-associated T cell.” These results show that CellReasoner generates expert-like reasoning chains and maps them into structured annotations, closing the loop from marker interpretation to decision-making. Its outputs reflect key expert traits: marker-by-marker integration, negative reasoning, contextual understanding, and decisive label assignment, validating the effectiveness of our CRAFT training strategy.

In conclusion, CellReasoner demonstrates that lightweight, open-source LLMs can be effectively adapted for cell type annotation using minimal supervision and carefully designed reasoning scaffolds. We benchmarked its inference efficiency on a single NVIDIA RTX 4090 GPU and found that inference time scaled approximately linearly with cell count, while maintaining a modest growth rate (**Supplementary Fig. 2e**), indicating that CellReasoner is suitable for large-scale datasets with consumer-grade hardware. Our CRAFT training strategy enables efficient knowledge activation from a small set of exemplars, delivering high accuracy, interpretability, and adaptability across diverse biological contexts, including cross-tissue, cross-cancer, and cross-modality settings. This work highlights the promise of reasoning-augmented LLMs in biomedical research, with potential applications extending beyond cell annotation to broader areas of omics interpretation, hypothesis generation, and knowledge-guided discovery.

## Methods

### Generation of a Global HVG Ranking

We collected a total of 570 single-cell studies from the curated cancer cell atlas ^18^ and TISCH2^17^ databases, spanning diverse tissues and disease contexts. For each dataset, we used the Seurat package to identify the top 3,000 HVGs, using gene symbols to represent their expression features. We then aggregated the HVG lists across all studies and computed the frequency of each gene’s occurrence across datasets. Genes were ranked in descending order based on their recurrence, resulting in a globally ordered list of HVGs across datasets (**Fig. 1a** and **Supplementary Table 2**). This list was subsequently used for constructing cell sentences and defining a unified input space for downstream tasks.

### Cell Sentence Construction

To standardize the model input, we selected the top *k* genes from the globally ranked HVG list as a unified reference feature space. For each individual cell *i*, we identified the subset of genes from this list that were expressed (i.e., non-zero expression), denoted as gene words 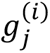. These gene words were retained in their original order as defined by the global HVG ranking and concatenated to form a natural language-style sequence, referred to as a cell sentence *C_i_* (Fig. 1b):

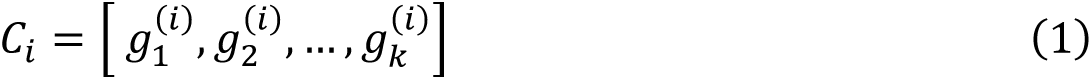

where *k* is a predefined maximum input length.

This approach ensures input consistency across datasets while preserving cell-specific expression features, thereby enabling the language model to perform cell-level reasoning.

### Problem Definition

To enable the model to integrate structured output with language-based reasoning, we designed it not only to generate cell type labels directly, but also to provide marker-by-marker reasoning chains that render the annotation process transparent and interpretable. We formulate single-cell type annotation as a task defined by a query 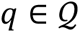, where 𝒬 denotes the space of annotation problems. Each query encapsulates the gene-level expression context of a single cell, represented as a cell sentence *C_i_* composed of gene words 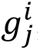, which are derived from the cell’s non-zero expressed genes. The objective is to generate both an annotation answer 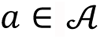 and a reasoning chain 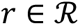. The reasoning chain *r* consists of a sequence of logical steps {*s*_1_,*s*_2_,…,*s_n_*}, where each step *s_i_* represents an intermediate inference from the input gene information to the final cell type annotation.

This formulation elevates cell type annotation from a label prediction task to a multi-step reasoning process that integrates cell-level gene expression profiles with known marker knowledge, thereby enhancing both accuracy and interpretability. Formally, the reasoning process is modeled as a function:

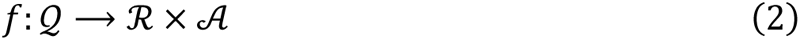

where 𝒬 denotes the space of annotation problems, 𝓡 is the space of reasoning steps; and 𝒜 is the set of possible cell type labels.

In addition, we define a simplified version of the task as:

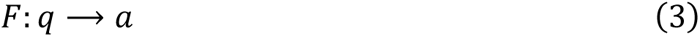

which directly maps a query *q* to an annotation label *a*.

### Reasoning Chain Construction

Chain-of-thought (CoT) language materials plays a critical role in training large language models, as it directly impacts their reasoning capability and overall performance. To ensure the accuracy and coherence of the reasoning paths, we employed a hybrid approach that combines AI-generated and human-curated expertise. Specifically, we first used a state-of-the-art reasoning model, DeepSeek-R1-671B, to generate initial CoT sequences. These preliminary reasoning chains were then manually reviewed, revised, and refined by human experts to enhance their logical rigor and interpretability. This process yielded a set of high-quality, validated reasoning chains for downstream model training.

### Dataset Construction

To support single-cell annotation, we designed two distinct task formulations, each guided by a specific instruction to steer the model’s reasoning behavior.

Task 1 – Direct Annotation Task

The first task instructs the model to infer the cell type directly based on marker gene expression: Instruction 1: *“These are highly expressed genes within a certain type of tumor cells; please annotate the cell type based on these marker genes. Please infer the cell type based on the marker genes, and directly give the final annotation result.”*

In this task, the model is expected to generate a cell type label directly from the input cell sentence containing highly expressed genes. The task is formally defined as:

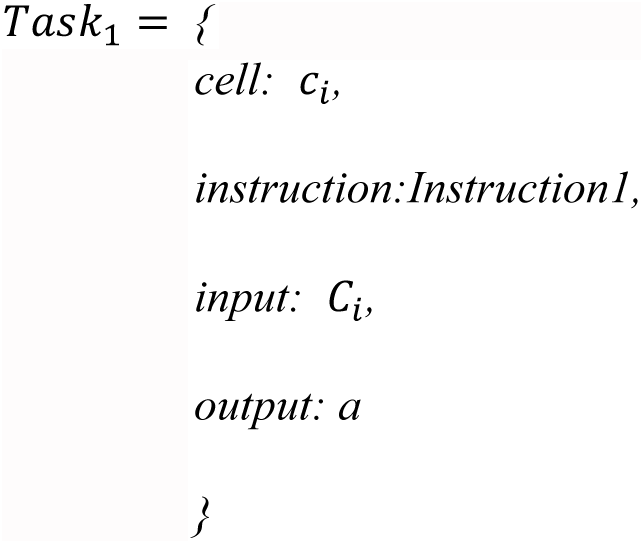

where *C_i_* denotes the cell sentence for cell *C_i_*, and 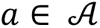 is the predicted cell type label. This task assesses the model’s ability to map cell sentence directly to a structured annotation output.

Task 2 – Reasoning-Augmented Annotation Task

The second task extends the first by requiring an explicit reasoning chain enclosed within specific tags:

Instruction 2: *“These are highly expressed genes within a certain type of tumor cells; please annotate the cell type based on these marker genes. Please infer the cell type based on the marker genes, and place the reason within the <*think*= and <*think*< tag.”*

In this version, the model is instructed not only to predict the cell type but also to provide a step-by-step reasoning process. The reasoning is embedded between <*think*= and <*think*< tags to ensure transparency and traceability. The task is defined as:

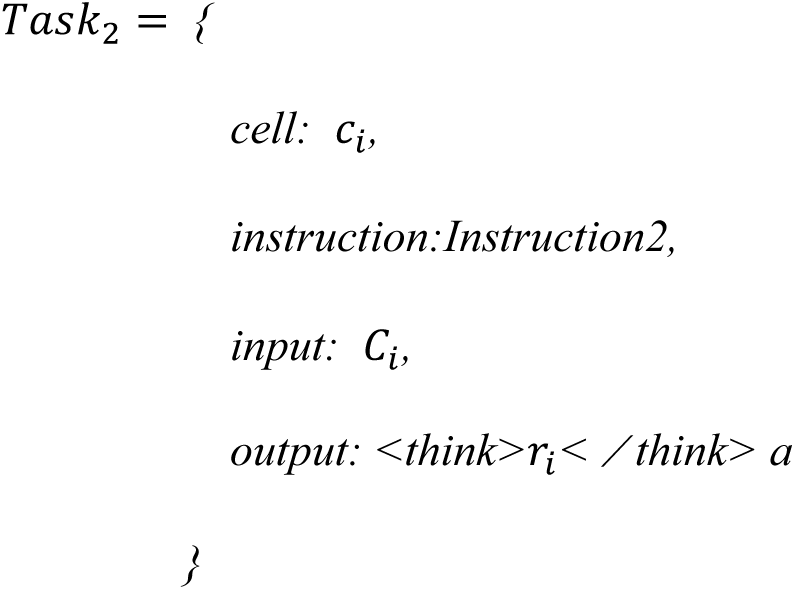

where 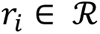 denotes the reasoning chain leading to the final annotation *a*. This design encourages interpretable output by requiring the model to articulate the logic behind its decision.

Together, these two instructional variants allow for the evaluation of model performance under different annotation paradigms—ranging from direct classification to reasoning-aware, explainable predictions.

### Training Workflow

#### CRAFT: Cell Reasoning and Annotation Fusion Training

Our experiments demonstrate that even small-scale LLMs possess a rich reservoir of latent biological knowledge within their parameter space (Supplementary Fig. 1a,b). This suggests that foundational biological concepts are already partially established during pretraining. However, such knowledge is often fragmented, forming isolated “knowledge islands” with only sparse and implicit connections between them.

We argue that the central challenge is no longer knowledge acquisition per se, but rather the activation of implicit knowledge through the construction of logical and interpretable connections. This shift—from knowledge learning to knowledge activation—is essential for effective biological reasoning (Fig. 1c). To address this challenge, we introduce CRAFT (Cell Reasoning and Annotation Fusion Training), a novel three-stage training framework designed to systematically activate the reasoning and annotation capabilities of small-parameter LLMs. CRAFT integrates carefully designed reasoning tasks with staged supervision to guide the model in constructing biologically coherent reasoning structures. This approach enhances both the effectiveness and interpretability of the model in single-cell annotation tasks.

Building on this insight, CellReasoner acquires its capabilities through a tightly coupled, three-stage CRAFT regimen:

- reasoning scaffold establishes initial, interpretable chains of thought using a minimal set of high-quality CoT exemplars.
- knowledge infusion enriches these nascent scaffolds by integrating a large corpus of biology-focused question–answer pairs, effectively bridging isolated knowledge islands.
- reasoning mode fusion realigns and fine-tunes the model’s inference patterns against CoT references, restoring latent reasoning pathways and further enhancing interpretability.

#### Stage I: Reasoning Scaffold

In this stage, we provide the model with an explicit reasoning framework by constructing preliminary reasoning chains based on cell types and their corresponding cell sentences. A total of 380 carefully curated, high-quality chain-of-thought samples are used for supervised training. During this phase, the model performs reasoning solely based on its prior knowledge, using the input gene sets as its sole evidence to infer the most likely cell type. The primary goal of this stage is to activate the model’s reasoning capability and to teach it how to associate gene sets with specific cell types in a step-by-step, expert-like manner. The resulting structure emulates domain-expert reasoning paths.

Formally, this process is modeled as:

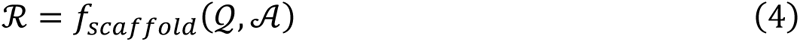

#### Stage II: Knowledge Infusion

In the second stage, we integrate more detailed biological knowledge into the model’s reasoning process. Through knowledge infusion, we activate previously isolated biological knowledge structures within the model, bridging these knowledge islands and creating a more cohesive and complete knowledge network. This infusion enhances the model’s internal representation of biological concepts, allowing it to incorporate additional relevant cell-type-specific knowledge and gene-to-gene relationships. As a result, the model’s cell type annotations become more accurate.

The goal of this stage is to enrich the model’s biological understanding, enabling it to make more precise inferences when faced with new data. However, after substantial knowledge infusion, we observe a decline in the model’s reasoning ability. This may be attributed to information overload or the increased complexity of integrating diverse knowledge, which can hinder the model’s capacity to effectively process all the infused knowledge and impact the precision of its reasoning.

We define the knowledge graph (*KG*) as the structured representation of this biological knowledge, and the process of constructing the knowledge network after infusion is modeled as:

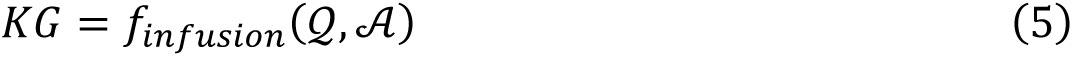

#### Stage III: Reasoning Mode Fusion

The third stage addresses the decline in reasoning ability observed after the knowledge infusion phase. To counteract this issue, we reintroduce the 380 high-quality chain-of-thought samples from the first stage for further training, fine-tuning the model to effectively reason and generalize within the knowledge network formed by the knowledge infusion. This approach enables the model not only to handle the newly injected biological knowledge more effectively, but also to integrate this knowledge with the pre-existing reasoning chains, ensuring that the reasoning process is both more efficient and precise. The core objective of this stage is to enable the model to identify optimal reasoning paths within complex knowledge networks, thereby restoring and enhancing its reasoning capacity and improving its overall performance in cell-type annotation tasks. The fusion model is thus capable of executing both complex reasoning and structured annotation, ensuring transparency and interpretability in the annotation process.

The optimization of the final reasoning path can be formally represented as:

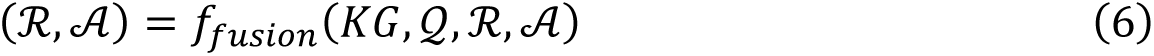

Through the three-stage training process and the optimization of the reasoning path, the model successfully transitions from knowledge learning to knowledge activation, evolving from isolated knowledge islands into a tightly integrated knowledge network. This transition ensures the efficiency, accuracy, and interpretability of the cell-type annotation task.

#### Loss Function

In the training of CellReasoner, we utilize the cross-entropy loss to measure the discrepancy between the model’s predicted probability distribution and the true label distribution. This loss function is used to update the model’s parameters, where *x_i_* represents the *i*-th input context (i.e., the cell sentence for a given cell), and *y_i_* is the corresponding target word (the next token).

The model’s predicted probability distribution is denoted *asp*(*y_i_*|*x_i_*). The cross-entropy loss is defined as:

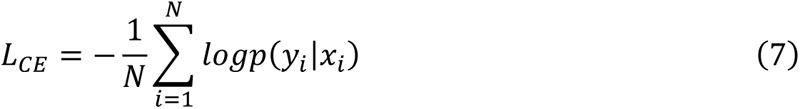

where *N* is the total number of samples. This loss function encourages the model to maximize the predicted probability corresponding to the true label at each forward pass, thereby continuously adjusting its parameters to bring the output distribution closer to the target distribution.

#### Training Dataset

We curated expert-annotated single-cell transcriptomic data from multiple high-quality pan-cancer datasets, encompassing a wide range of tumor and immune cell types. In total, 38 distinct cell types were included (**Supplementary Table 3**). To support model training and evaluation, we constructed multiple datasets following the strategy below:

First, to support reasoning ability learning, we constructed a high-quality CoT dataset by randomly sampling 10 cells from each of the 38 cell types, resulting in 380 carefully selected samples (**Supplementary Table 4**). During the sampling process, we enforced a constraint that the gene-word overlap between any two cell sentences must not exceed 50%, in order to enhance sample diversity and improve reasoning generalizability.

Second, to support training in the knowledge infusion stage, we randomly selected 1,000 cells per cell type, resulting in a pancancer38k training dataset consisting of 37,187 cells (**Supplementary Table 5**). For cell types with fewer than 1,000 available cells, we retained 100 cells for internal evaluation and used the remainder for training.

Finally, we randomly selected an additional 3,800 cells to construct an internal test set (**Supplementary Table 6**), ensuring that there was no overlap with the training data. This dataset was used for downstream model evaluation and analysis. For all training and test sets, we used the top 1,500 globally ranked HVGs as input features.

#### Implementation Details

We used format-specific tools to extract gene lists from single-cell datasets based on their file types. For datasets in rds format, we employed the Seurat package for data extraction and processing, while for datasets in h5ad format, we used Scanpy to retrieve gene information.

Throughout all three stages of CRAFT training, we applied supervised fine-tuning (SFT) with cross-entropy loss to refine the model’s predictions at each stage.. During both the reasoning scaffold and reasoning mode fusion stages, training was conducted exclusively on 380 high-quality CoT samples. Subsequently, the knowledge infusion stage leveraged 37,187 question– answer pairs to reinforce the model’s internal knowledge representation.

We used consistent hyperparameters across all models within the same stage, with each trained on a single A800 GPU. Model training was carried out using the Llama-factory framework with Low-Rank Adaptation (LoRA) ^19^ for parameter-efficient fine-tuning. We used hyperparameters with a LoRA matrix rank of 8 and a scaling factor of 16. The batch size was set to 8 for the reasoning scaffold and reasoning mode fusion stages, and 20 for the knowledge infusion stage. The learning rate was uniformly set to 5e-5, and the maximum input sequence length was 8,192 tokens. To ensure stable and efficient training, we employed a cosine learning rate scheduler.

During inference, we adopted task-specific decoding strategies to balance generation stability and reasoning quality. For direct annotation tasks (Task 1), we employed greedy decoding by selecting the most probable token at each step to ensure deterministic outputs. For reasoning-intensive tasks (Task 2), we applied nucleus sampling with a top-p value of 0.4 and a temperature of 0.6 to enhance the diversity and coherence of the generated chain-of-thought responses.

### Benchmarking Methods

#### ChatGPT -4o, ChatGPT o3, DeepSeek-V3, and DeepSeek-R1

To ensure fairness in our comparative analysis, we standardized the annotation procedures across all evaluated models. For the cluster-marker-based annotation task, ChatGPT -4o (April 29, 2025 version), ChatGPT o3 (April 16, 2024 version), DeepSeek-V3 (March 25, 2025 version), and DeepSeek-R1 (January 20, 2025 version) followed an identical pipeline. Specifically, cells were first clustered using the Louvain algorithm with a resolution parameter set to 0.5. From each resulting cluster, the top-k marker genes were extracted and provided as input to each model. All models received the same prompt, consistent with our Task 1 instruction: to directly predict the cell type label based on the provided marker genes. In terms of execution, ChatGPT -4o and ChatGPT o3 were accessed via the OpenAI official web interface while DeepSeek-V3 and DeepSeek-R1 were evaluated through API-based submissions. Additionally, for experiments involving cell sentences constructed from top-ranked HVGs, only DeepSeek-V3 and DeepSeek-R1 were evaluated under this setting.

#### SingleR

We additionally employed SingleR^20^ (version 1.4.1) as a conventional reference-based annotation method. SingleR determines the most likely cell type for each cell by comparing its expression profile with reference expression patterns. In this study, we used the BlueprintEncodeData dataset from the celldex package as the reference, which includes expression profiles for a broad range of human immune cell types. Annotation was performed at the single-cell level, with labels assigned based on similarity scores.

#### Evaluation of Cell Type Annotations

To assess the accuracy of predicted cell type labels, we designed an evaluation framework that combines automated scoring using a large language model (LLM) with manual verification. Specifically, we used DeepSeek-V3 to assess the semantic consistency between the reference label (A) and the predicted label (B).

The evaluation is based on a structured prompt that defines four levels of label consistency:

- Subtype correct: A and B refer to the same cell subtype, possibly using different naming conventions; alternatively, B is a more specific subtype of A.
- Major correct: A and B belong to the same major cell category but represent different subtypes; alternatively, A is a more specific subtype of B.
- Partially correct: B includes multiple potential labels, some of which match A (either as Subtype or Major correct), but others are incorrect or irrelevant, resulting in ambiguity.
- Incorrect: A and B belong to entirely different cell lineages, or B includes multiple predictions with none matching A.

The exact prompt used for automated evaluation is provided below:

*“I am working on a single-cell transcriptomic cell type annotation task. The reference label (A) may be an abbreviation or a marker-based description of a cell type. Both the reference label(A) and the model’s predicted label (B) have been standardized and mapped to Cell Ontology terms whenever possible. As an expert in single-cell annotation, please evaluate whether my model’s predicted label (B) is accurate, according to the following criteria:*

- *Major correct: A and B belong to the same major cell type category (e.g., T cells), but are different subtypes (e.g., CD4+ vs CD8+), or A is a subtype of B, and B is the major category of A (e.g., A = CD4+ T cells, B = T cells);*
- *Subtype correct: A and B refer to the same cell subtype, despite differences in naming conventions (e.g., abbreviation vs marker gene expression), or B is a subtype of A (e.g., A = B cells, B = class-switched memory B cells);*
- *Partially correct: B contains multiple possible cell types (e.g., “celltypeA or celltypeB”), and one of them matches A (either Major or Subtype correct), but others are incorrect or unrelated, making the prediction ambiguous;*
- *Incorrect: B and A belong to entirely different cell lineages (e.g., labeling a neuron as an epithelial cell), or B includes multiple predictions and none of them match A;*

*Please follow the output format below:*

- *First, provide a brief reasoning explaining your judgment*.
- *End with a line formatted exactly as Judgment: [Category], where the category must be one of: Major correct / Subtype correct / Partially correct / Incorrect*.
- *Example output:* *Reasoning: A and B are both T cell types, but A is CD4+ and B is CD8+, indicating different subtypes within the same major category*. *Judgment: Major correct*

*Now evaluate the following case:”*

All LLM-generated responses were manually reviewed and corrected when necessary to ensure consistency and reliability across test cases. For quantitative comparison, we assigned numerical scores to each evaluation category: Subtype correct = 1.0, Major correct = 0.5, Partially correct = 0.25, and Incorrect = 0. The complete set of paired evaluations between model predictions and ground-truth labels is provided in Supplementary Table 7.

### Datasets PDAC dataset

The PDAC dataset (Pancreatic Ductal Adenocarcinoma) comprises 57,702 single-cell transcriptomes encompassing 51 distinct cell subtypes. It includes a broad spectrum of immune and non-immune cell types, such as CD8+ T cells, NKT cells, CD4+ T cells, plasma cells, monocytes, neutrophils, fibroblasts, epithelial cells (EC), endothelial cells, and B cells. The dataset also contains various macrophage and T cell subsets, endocrine cells, acinar cells, T follicular helper (TFH) cells, and mast cells. Notably, macrophage subtypes include LAM1–4, M1-like, and M2-like macrophages.

### PBMC3K dataset

The PBMC3K dataset contains single-cell RNA-seq data from 2,700 peripheral blood mononuclear cells (PBMCs) derived from a healthy human donor. The dataset includes major immune cell populations such as naïve CD4+ T cells, memory CD4+ T cells, CD14+ monocytes, B cells, CD8+ T cells, FCGR3A+ monocytes, natural killer (NK) cells, dendritic cells (DCs), and platelets.

### Multiome PBMC dataset

This dataset contains paired single-cell RNA and ATAC-seq profiles from human PBMCs, generated using the 10x Genomics Single Cell Multiome platform. After quality control filtering, the dataset comprises 10,412 cells annotated into 19 cell types, including CD14+ monocytes, CD16+ monocytes, CD4+ naive T cells, CD4+ central memory (TCM) and effector memory (TEM) T cells, CD8+ naïve and TEM subsets, conventional dendritic cells (cDCs), γδ T cells, hematopoietic stem and progenitor cells (HSPCs), intermediate and memory B cells, naïve B cells, NK cells, plasmacytoid DCs (pDCs), plasma cells, mucosal-associated invariant T (MAIT) cells, and regulatory T cells (Tregs).

### Liver dataset

The liver dataset comprises 8,444 single-cell transcriptomes from human liver tissue. Cell types include αβ T cells, γδ T cells, NK-like cells, mature B cells, plasma cells, erythroid cells, cholangiocytes, hepatocytes, hepatic stellate cells, inflammatory and non-inflammatory macrophages, periportal and central venous liver sinusoidal endothelial cells (LSECs), and portal endothelial cells.

## Data availability

All data used in this manuscript are publicly available and the sources are fully cited. Specifically, the PBMC3K dataset was obtained from https://www.10xgenomics.com/cn/datasets/3-k-pbm-cs-from-a-healthy-donor-1-standard-1-1-0, and the Multiome PBMC dataset was downloaded from https://support.10xgenomics.com/single-cell-multiome-atac-gex/datasets/1.0.0/pbmc_granulocyte_sorted_10k. The liver dataset was downloaded from https://www.ncbi.nlm.nih.gov/geo/query/acc.cgi?acc=GSE115469. The PDAC dataset was obtained from https://www.ncbi.nlm.nih.gov/geo/query/acc.cgi?acc=GSE197177.

## Code availability

The codebase for CellReasoner is publicly available at https://github.com/compbioNJU/CellReasoner. The models can be accessed at the following Hugging Face links: CellReasoner-7B: https://huggingface.co/guangshuo/CellReasoner-7B, CellReasoner-32B: https://huggingface.co/guangshuo/CellReasoner-32B, CellReasoner-liver-10-shot-7B: https://huggingface.co/guangshuo/CellReasoner-liver-10-shot-7B, and CellReasoner-scATAC-10-shot-7B: https://huggingface.co/guangshuo/CellReasoner-scATAC-10-shot-7B

## Acknowledgements

The authors acknowledge the High Performance Computing Center of Nanjing University for providing high performance computing (HPC) resources. This work is supported by the Postgraduate Research & Practice Innovation Program of Jiangsu Province.

## Author contributions

D.C. and M.C. supervised the study. G.C. designed the study, implemented the model and performed data analysis with support from Y.S. and J.W.. H.C. assisted with data processing.

G.C. wrote the manuscript with input from Y.S.. All authors reviewed and approved the submitted manuscript.

