## Supplementary Figure1-3 for "CellReasoner: A reasoning-enhanced large language model for cell type annotation"

*by* Cao *el al.*

**a** You are a gene-function annotation expert. When I provide a gene or a list of genes (gene set), please automatically produce a table with the following columns:
• Gene • Basic description (e.g. protein family or primary function) • Pathways or molecular mechanisms involved • Known roles in diseases or physiological processes
Input: CD4, IL7R, CD40LG,CD69

| Gene | Basic Description | Pathways or Molecular Mechanisms | Known Roles in Diseases or Physiological Processes |
|---|---|---|---|
| CD4 | T-cell surface protein (CD4) that is a coreceptor for the HIV-1 envelope protein | T-cell activation, HIV-1 infection | AIDS, T-cell lymphomas, autoimmune diseases, HIV-1 infection |
| IL7R | Interleukin-7 receptor, a type I cytokine receptor | T-cell development, B-cell development, hematopoiesis | Severe combined immunodeficiency (SCID), autoimmune diseases, lymphomas |
| CD40LG | CD40 ligand, a member of the tumor necrosis factor (TNF) superfamily | B-cell activation, T-cell activation, immune response | Multiple myeloma, lymphomas, autoimmune diseases, inflammatory diseases |
| CD69 | Early activation marker on T cells, B cells, and NK cells | T-cell activation, B-cell activation, NK-cell activation | Autoimmune diseases, inflammatory diseases, viral infections |

**b** You are an expert in cell biology and gene markers. When I provide a cell type, please automatically generate a table with the following columns:
• Marker gene • Function or biological significance • Expression characteristics in this cell type
Input: Cell type: CD4 T cell

| Marker Gene | Function or Biological Significance | Expression Characteristics in CD4 T Cells |
|---|---|---|
| CD4 | T-cell surface protein that is a coreceptor for the HIV-1 envelope protein | High expression, essential for CD4 T cell identity and function |
| CD28 | T-cell receptor-associated protein that provides co-stimulation signals | High expression, essential for T-cell activation and proliferation |
| CD45RA | T-cell receptor (TCR) associated protein that is a marker for naive T cells | High expression in naive CD4 T cells, downregulated in memory CD4 T cells |
| CD45RO | T-cell receptor (TCR) associated protein that is a marker for memory T cells | High expression in memory CD4 T cells, downregulated in naive CD4 T cells |
| CCR7 | Chemokine receptor that is involved in T-cell homing to lymphoid tissues | High expression in naive CD4 T cells, downregulated in memory CD4 T cells |
| CD69 | Early activation marker that is upregulated in activated T cells | High expression in activated CD4 T cells, downregulated in resting cells |

**c** GPR183, LYZ, GZMA, PRF1, GZMK, LTB, SERPINA1, NKG7, CD69, IL7R, CCL5, ALOX5AP, CCL20, CD8A,…, CPPED1, MAP4K4, PPP1CB, RPL36, RPL7A, RPS15A, SEC61B, SLC18B1.
**These are highly expressed genes within a certain type of tumor cells; please annotate the cell type based on these marker genes. Please infer the cell type based on the marker genes, and directly give the final annotation result.**

CellReasoner-7B: CD8

**d** GPR183, LYZ, CD74, GZMA, MT2A, HLA-DQB1, LTB, CD69, IL7R,, CCL5, DUSP2, ALOX5AP, TIMP1, ANXA1, ITM2A, SPON2, LGALS1, ..., RPL7A, RPS15A, SH3BGRL, SLC25A3.
**These are highly expressed genes within a certain type of tumor cells; please annotate the cell type based on these marker genes. Please infer the cell type based on the marker genes, and place the reason within the <think></think> tag.**

**Qwen2.5-7B-Instruct base** (✗)

Based on the provided marker genes, this list predominantly includes a mix of immune-related and tumor-associated genes. However, a significant portion of these genes are known to be associated with T cells, which are a type of white blood cell that plays a central role in the immune system. Notable genes like CD3D, CD3G, CD37, CD4, CD52, CD2, CD40LG, CD48, CD7, CD96, CD99, CD247, CD44, CD59, CD63, CD69, CD74, CD8A, CD8B, CD96, CD147, CD154, CD160, CD161, CD166, CD178, CD19, CD27, CD28, CD30, CD38, CD40, CD45, CD45RA, CD45RO, CD45RO, CD57, CD58, CD69, CD80, CD86, CD117, CD123, CD137, CD152, CD169, CD195, CD197, CD229, CD252, CD274, CD278, CD289, CD335, CD361, CD37, CD38, CD39, CD40, CD40L, CD45, CD45RA, CD45RO, CD45RO, CD57, CD58, CD69, CD80, CD86, CD117, CD123, CD137, CD152, CD169, CD195, CD197, CD229, CD252, CD274, CD278, CD289, CD335, CD361, CD37, CD38, CD39, CD40, CD40L, CD45, CD45RA, CD45RO, CD45RO, CD57, CD58, CD69, CD80, CD86, CD117, CD123, CD137, CD152, CD169, CD195, CD197, CD229, CD252, CD274, CD278, CD289, CD335, CD361, CD37, CD38, CD39, CD40, CD40L, CD45, CD45RA, CD45RO, CD45RO, CD57, CD58, CD69, CD80, CD86, CD117, CD123, CD137, CD152, CD169, CD195, CD197, CD229, CD252, CD274, CD278, CD289, CD335, CD361, CD37, CD38, CD39, CD40, CD40L, CD45, CD45RA, CD45RO, CD45RO, CD57, CD58, CD69, CD80, CD86, CD117, CD123, CD137, CD152, CD169, CD195, CD197, CD229, CD252, CD274, CD278, CD289, CD335, CD361, CD37, CD38, CD39, CD40, CD40L, CD45, CD45RA, CD45RO, CD45RO, CD57, CD58, CD69, …

meaningless, repetitive CD-marker dump; lacks substantive analysis; no effective answer.

overextended conclusion

ambiguous label

**DeepSeek-R1**

<think> Okay, let's tackle this problem. The user provided a long list of genes that are highly expressed in a certain type of tumor cells and wants me to annotate the cell type based on these markers.

First, I need to recall common cell types and their associated marker genes. Looking through the list, I notice several genes related to T cells. **For example, CD3D, CD3G, and CD247** are part of the CD3 complex, which is specific to T cells. Also, CD2, CD7, CD4, and CD8 (though CD8 isn't listed here) are typical T cell markers.

There's also **IL7R (Interleukin-7 receptor), which is important for T cell development and memory**. SELL (L-selectin) is found on naive T cells. Genes like GZMA (Granzyme A) and CCL5 are involved in cytotoxic functions, suggesting cytotoxic T cells. Additionally, genes like LTB (lymphotoxin beta) and CD69 (an activation marker) are associated with T cell activity.

**Other markers like** CD52, CD48, and LCK (a kinase involved in T cell receptor signaling) further support the T cell hypothesis. The presence of genes like CXCR4 and S1PR1 might relate to T cell trafficking and migration.

**Looking at the list again**, there's no strong indication of B cell markers (like CD19, CD20) or myeloid markers (like CD14, CD68). Some genes like HLA molecules are present, but those are more general. The combination of T cell receptor components, activation markers, and cytotoxic molecules points towards T cells, specifically possibly activated or tumor-infiltrating T cells.

**Given the tumor context, these could be tumor-infiltrating lymphocytes (TILs), which are T cells that have left the bloodstream and migrated into a tumor.** However, since the question is about the cell type of the tumor cells, if these markers are expressed by the tumor cells, it might indicate a T cell lymphoma. **But typically, T cell lymphomas would express T cell markers. Alternatively, if the tumor is of a different origin but has a microenvironment rich in T cells, but the question states ""highly expressed genes within a certain type of tumor cells,"" implying the tumor cells themselves express these.**

Wait, but tumors like T-cell acute lymphoblastic leukemia (T-ALL) or T-cell lymphomas would express T cell markers. Alternatively, maybe the tumor is derived from T cells. The presence of multiple T cell-specific genes (CD3D, CD3G, CD247, TRAT1, CD2, etc.) strongly suggests that the tumor cells are of T cell origin. **Therefore, the cell type is likely T cells, and the tumor type would be a T cell lymphoma or leukemia.**

So, the final annotation based on these markers is T cells, leading to the conclusion that the tumor cells are of T cell origin, **hence a T cell lymphoma or leukemia**. The key markers here are the CD3 complex genes and other T cell-specific genes. </think>

Annotation: **T cells (likely T cell lymphoma or tumor-infiltrating T lymphocytes)**

marker-by-marker reasoning

lack of focused inference path, oscillating assumptions

**CellReasoner-7B** (✓)

<think>First, I need to recall what each gene is typically associated with.

CD3D and CD3G are part of the T-cell receptor complex, so that's a strong indicator of T-cells. CD4 is another T-cell marker, especially for helper T-cells. Then there's CD2, which is common in lymphocytes like T-cells and B-cells. CD7 is also found on T-cells. **The presence of IL7R (interleukin-7 receptor) suggests activated or memory T-cells since they express this after activation.**

**Looking at other markers**: LYZ encodes lysozyme, which might be present in myeloid cells but can also be in some T-cell subsets. GZMA is granzyme A, often associated with cytotoxic cells like NK cells or CD8+ T-cells. However, some CD4+ T-cell subsets can express granzymes under certain conditions, especially when activated.

**HLA-DQB1** is involved in antigen presentation, which T-cells use to present antigens to other immune cells, supporting an adaptive immune cell identity. CD69 is an early activation marker on T-cells. IL32 is secreted by activated T-cells. CCL5 (RANTES) is a chemokine produced by T-cells during inflammation.

**CXCR4** is a chemokine receptor important for homing T-cells to specific tissues. SELL (L-selectin) is involved in initial T-cell binding to endothelial cells. CD48 is a molecule involved in T-cell activation and regulation. CD52 is targeted by alemtuzumab, used in treating autoimmune diseases and leukemia, and is found on lymphocytes including T-cells.

**Other genes like FOS and JUN** are immediate early genes involved in various cellular responses, including T-cell activation. Genes such as TNFRSF1B (CD120b) are involved in apoptosis and cytokine signaling in T-cells.

**Some genes like VIM (vimentin)** are cytoskeletal proteins not specific to T-cells but might indicate active cells. TIMP1 is involved in extracellular matrix remodeling, possibly related to T-cell migration.

**Putting this together,** the majority of the markers point towards T-cells due to CD3, CD4, IL7R, and others. **The presence of granzymes and cytotoxic molecules could suggest a subset of CD4+ T-cells with effector functions**, though some literature indicates CD4+ T-cells can express granzymes upon activation. Additionally, HLA class II genes like HLA-DQB1 support antigen-presenting capabilities typical of T-helper cells.

**There's no strong evidence here for** B-cells (CD19, CD20 missing), plasma cells (CD138), or myeloid cells (like CD14, CD68). NK cells would have different markers like CD56 (NCAM1) and CD16 (FCGR3A), which aren't listed here. **Therefore, the most likely cell type is CD4+ T-cells.**</think>

**CD4**

evidence-based synthesis, marker-to-function mapping

negative exclusion, deductive logic, cell-type focus

precise label

**Supplementary Figure 1. Cell level annotation and reasoning capabilities of CellReasoner.** **a**, Gene level biological knowledge retrieval by Qwen2.5-7B-Instruct, including protein function, pathway roles, and disease associations. **b**, Cell type–specific marker function retrieval in the context of CD4 T cells. **c**, Direct cell type annotation by CellReasoner-7B without reasoning output. **d**, Expert-level reasoning output by CellReasoner-7B, integrating marker functions (e.g., CD3, IL7R) to infer the final label. Compared to DeepSeek-R1 and Qwen2.5-7B-Instruct, CellReasoner provides more accurate and interpretable predictions.

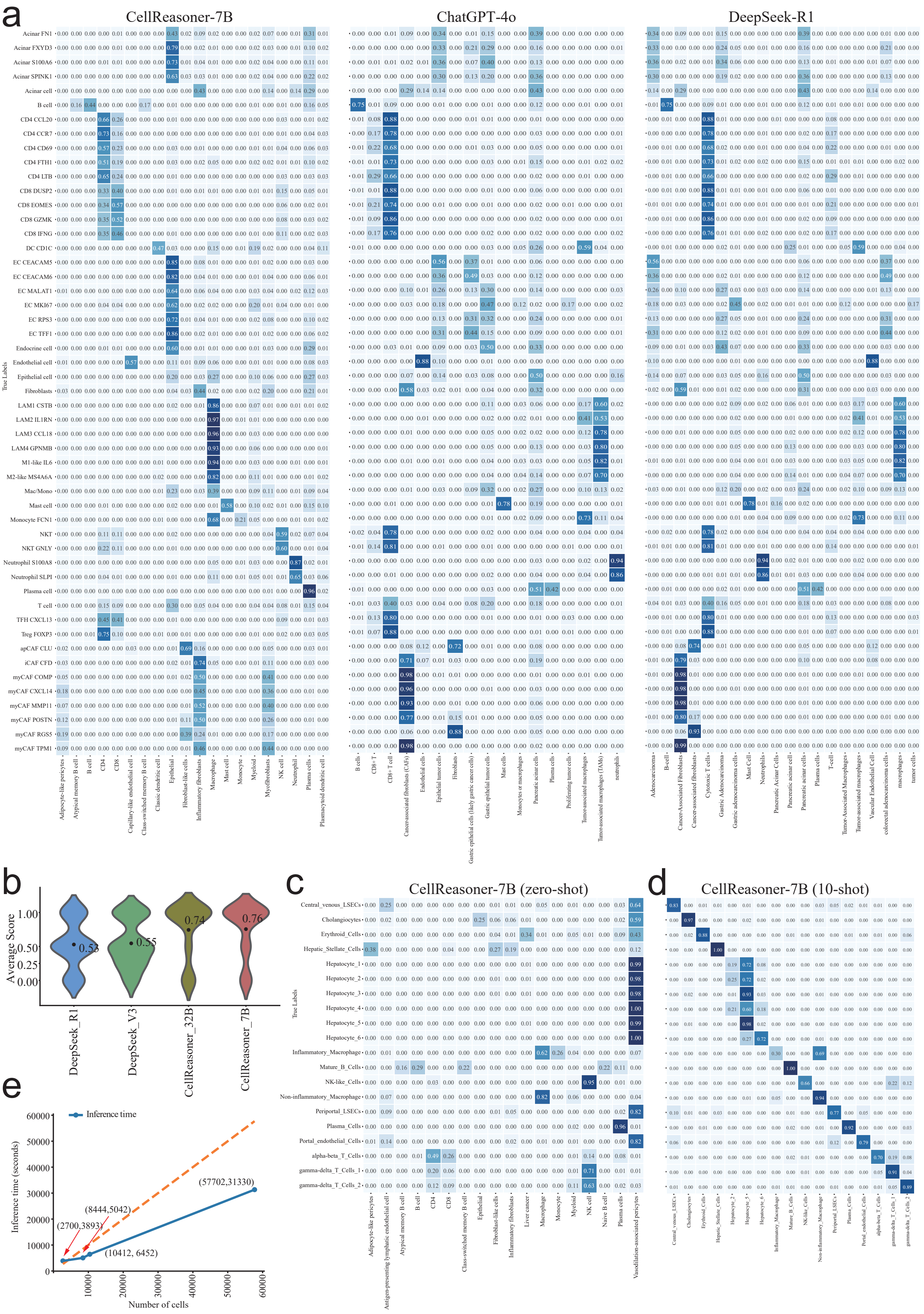
a
CellReasoner-7B
ChatGPT-4o
DeepSeek-R1
True Labels
b
Average Score
1.00
0.50
0.25
0.00
0.53
0.55
0.74
0.76
DeepSeek_R1
DeepSeek_V3
CellReasoner_32B
CellReasoner_7B
c
CellReasoner-7B (zero-shot)
d
CellReasoner-7B (10-shot)
e
Inference time
Inference time (seconds)
Number of cells
(2700,3893)
(8444,5042)
(10412, 6452)
(57702,31330)

**Supplementary Figure 2. CellReasoner performance and inference efficiency.** **a**, Confusion matrices on the PDAC dataset for CellReasoner-7B (top-1000 HVGs), ChatGPT-o4 (top-20 markers), and DeepSeek-R1 (Top-50 markers). **b**, Average scores on a class-balanced PDAC subset (20 cells per type). **c-d**, Zero-shot and 10-shot annotation on the liver dataset using CellReasoner-7B. **e**, Inference time evaluation on a single NVIDIA RTX 4090 GPU across datasets of increasing cell counts.

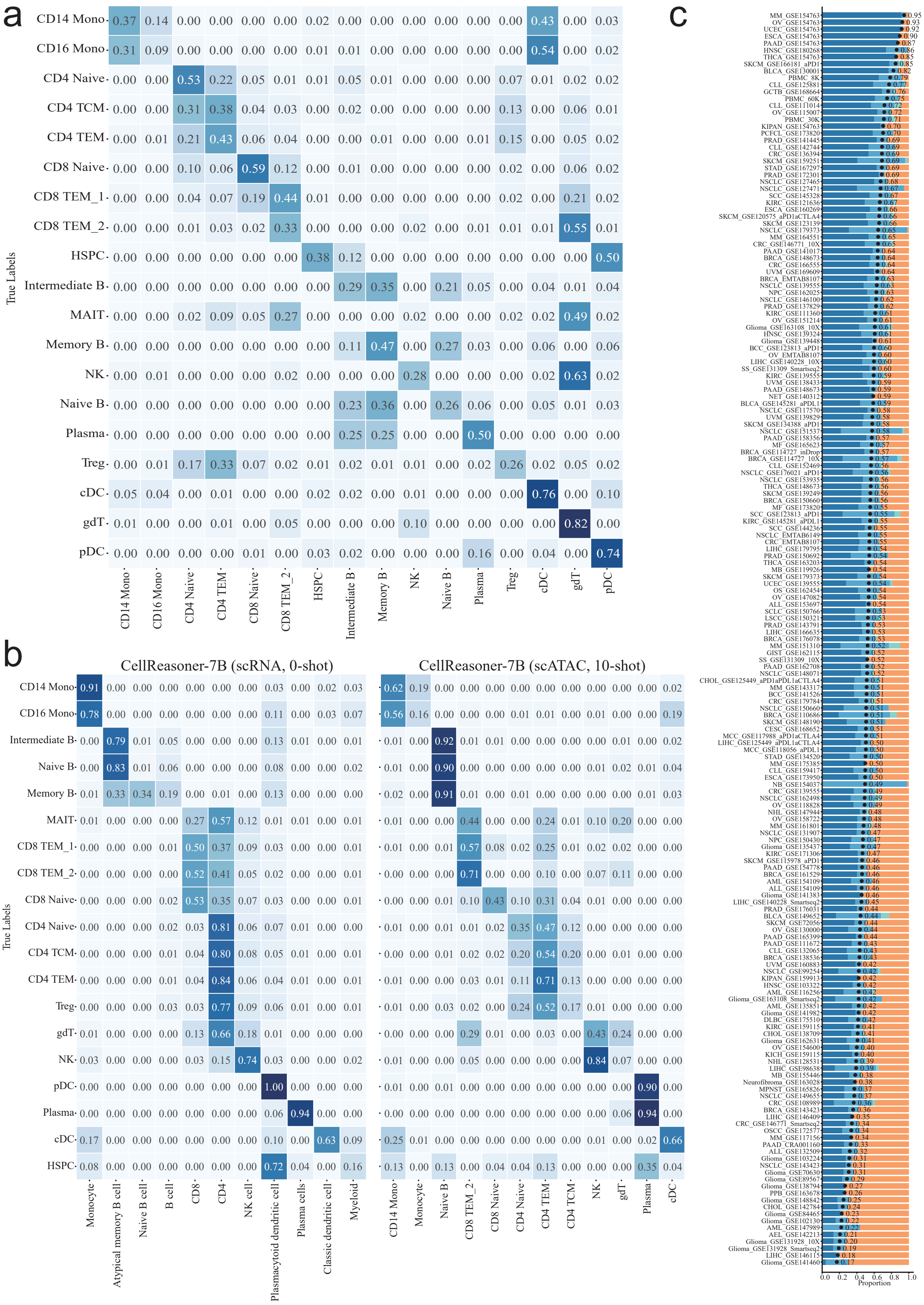
a
b
c
CellReasoner-7B (scRNA, 0-shot)
CellReasoner-7B (scATAC, 10-shot)
True Labels
Proportion
CD14 Mono
CD16 Mono
CD4 Naive
CD4 TCM
CD4 TEM
CD8 Naive
CD8 TEM_1
CD8 TEM_2
HSPC
Intermediate B
MAIT
Memory B
NK
Naive B
Plasma
Treg
cDC
gdT
pDC
Monocyte
Atypical memory B cell
Naive B cell
B cell
CD8
CD4
NK cell
Plasmacytoid dendritic cell
Plasma cells
Classic dendritic cell
Myeloid

**Supplementary Figure 3. Cross-modality and large-scale annotation performance of CellReasoner.** **a**, Confusion matrix of CellReasoner-7B on the multiome PBMC scATAC-seq dataset under 10-shot setting. **b**, Performance on the paired scRNA-seq modality of multiome PBMC. Left: zero-shot (no fine-tuning); Right: after 10-shot training on scATAC-seq. **c**, Zero-shot annotation performance across 181 datasets spanning 48 cancer types.
